# *Apolipoprotein E ε4* exacerbates microglia-mediated complement-dependent synapse loss caused by neuronal *Tpk* deficiency

**DOI:** 10.1101/2025.01.16.633468

**Authors:** Pengyue Du, Boru Jin, Zijie Wang, Binqiao Zhao, Shaoming Sang, Chunjiu Zhong

## Abstract

Thiamine pyrophosphokinse-1 (TPK) is a key enzyme that converts thiamine to functional thiamine diphosphate (TDP). TPK insufficiency and hence TDP reduction in neurons induced by amyloid-β deposition and diabetes, an independent risk factor of Alzheimer’s disease (AD), recapitulate multi-pathophysiological features in the brain of mice, similar to those in human AD. *Apolipoprotein E* ε*4 allele* (*APOE4*) is the most well-known genetic risk factor for AD. Clinical trials by boosting TDP using benfotiamine, a derivative of thiamine that significantly elevates TDP level in human encrythrocytes, have shown the inferior clinical efficacies in *APOE4* carriers compared to non-APOE4 carriers. Clarifying the relationship between APOE4 and TPK expression and multi-pathophysiological characteristics of AD induced by *Tpk* deficiency in neurons is imperative. Here, we find that humanized *APOE4* didn’t directly affect *Tpk* expression of mice, but markedly aggravates behavior abnormalities of *Tpk-*cKO mice. Pathologically, the *Tpk-*cKO mice with humanized *APOE4* knock-in (AE-cKO mice) exhibit more synapse loss than the mice with only humanized *APOE4* knock-in and the *Tpk-*cKO mice. Transcriptomics and pathologic analysis identified that *APOE4* promoted the overactivation of microglia and the transition of microglia to a disease-associated and phagocytosis state *via* a complement-mediated pathway. Further, the C3aR antagonist significantly repressed microglia phagocytosis and synaptic elimination of the AE-cKO mice. Our results demonstrate that *APOE4* exacerbates behavior dysfunction of *Tpk-*cKO mice through microglia-mediated complement-dependent synaptic elimination. These findings provide important insights into the role of APOE4 in the pathogenesis of AD.

## Introduction

Alzheimer’s disease (AD) is a complex neurodegenerative disease, which is involved in various risk factors and multiple pathophysiological features. The multiple pathophysiological features of AD include β-amyloid (Aβ) deposition forming plaques, glial activation and neuroinflammation, brain glucose hypometabolism, Tau hyperphosphorylation aggregating to neurofibrillary tangles, and progressive loss of synapses and neurons causing brain atrophy [1, 2]. Among those features, progressive loss of synapses and neurons causing brain atrophy have been recognized as the direct pathological event that underlies clinical symptoms. However, the mechanism(s) is not fully elucidated.

The risk factors of AD can be classified into aging, genetic factors, chronic diseases and pathological conditions, unhealthy lifestyles, and educational status [3]. Among them, the *Apolipoprotein E* ε*4* allele (*APOE4*) is the most well-known genetic risk factor for AD. Compared with non-*APOE4* (ε2 and ε3 alleles) carriers, heterozygous *APOE4* carriers show a two- to three-fold increased risk of AD and an average onset age of about 6 years earlier, while homozygous carriers exhibit ten- to twelve-fold increased risk of AD and an average onset age of about 14 years earlier [4, 5]. The mechanisms by which *APOE4* significantly increases the risk of AD have been widely explored, which are demonstrated to be involved in exacerbating Aβ pathology [6, 7], inducing Tau pathology [8–11], perturbing lipid and glucose metabolism, glial activation and neuroinflammation, and so forth [9, 12]. However, the mechanisms by which *APOE4* promotes the occurrence and development of AD are too much variations in different pathological conditions [9, 13, 14]. It is still inconclusive how *APOE4* leads to AD.

Our previous studies have demonstrated that thiamine pyrophosphokinase-1 (TPK), a key enzyme that converts thiamine into functional thiamine diphosphate (TDP), is specifically inhibited in brain samples from AD patients [15]. Further, conditional *Tpk* knockout in excitatory neurons of adult mice (*Tpk-*cKO) induces all important AD-associated pathophysiological features, including brain Aβ deposition and plaques, Tau hyperphosphorylation and neurofibrillary tangles, glial activation and neuroinflammation, and progressive loss of synapses and neurons causing brain atrophy [15, 16]. Furthermore, *Tpk* re-expression and thiamine/TDP supplement manifest significant rescue effects on pathophysiological alterations and behavioral abnormalities caused by *Tpk* insufficiency in both cellular and mouse models [16]. The results of two phase trials and an open clinical observation study of benfotiamine, a lipid-soluble thiamine derivative that significantly elevates the level of TDP in human blood and erythrocyte, exhibit powerfully beneficial effects in delaying the cognitive decline of AD patients [16–18]. Therefore, our explorations on the role of TPK inhibition and hence TDP reduction in the pathogenesis of AD provide innovative insights into the pathogenesis and treatment of the disease. More importantly, the two phase clinical trials of benfotiamine showed that the clinical efficacies of *APOE4* carriers were significantly inferior to those in non-carriers [16, 17]. Understanding the relationship between APOE4 and TPK expression and multi-pathological cascades induced by *Tpk* deletion in neurons will represent an opportunity for exploring therapeutic avenues of AD by targeting APOE4.

To address those issues, our current studies further investigate the effects of APOE4 on brain TPK expression and pathophysiological features caused by neuronal *Tpk* deletion in adult mice. The results show that APOE4 significantly worsens the abnormalities of behavior tests and the loss of synapse in the *Tpk-*cKO mice, which the mechanism is involved in the excessive pruning of synapses mediated by microglia-associated complement pathways.

## Methods

### Animals

All animal care and experimental procedures were approved by the Medical Experimental Animal Administrative Committee of Fudan University. The conditional knockout of m*Tpk* allele (NM_013861.3), which has nine exons with the ATG start codon in exon 2 and TAA stop codon in exon 9, exon 4 was selected for targeting. The deletion of exon 4 results in the loss of function of *Tpk* gene by a frameshift. The detailed procedures have been described in our previous study [15]. The mice with humanized *APOE4* knock-in (KI, NM-HU-190002) were purchased from Shanghai Model Organisms.

### Behavior tests Nesting test

The mice were placed in separate cages one hour before dark and added nesting material. The status of the mice and nesting materials was observed after12 hours of darkening. The nesting behavior was rated on a scale of 0-5, with 5 representing normal and 0 representing worst.

### Y-maze test

The Y-maze device consists of 3 black horizontal arms (36 cm long, 5 cm wide, and 10 cm high) at 120° angles to each other. The mouse was placed in the center of the maze and let it explore freely for 5 min, recording the rate of alternation of the mouse in the maze.

### Rotarod test

Rotarod test is used to test motor function of mice. The rod initially rotates at a constant speed of 4 rpm to allow all mice to be positioned in their respective channels. Once all the mice are on the bar, press the start button and the bar accelerates from 40 rpm to 300 rpm in 40 seconds. The rate of mice falling off the bar is recorded.

### Western blotting

Brain tissues were lysed in RIPA lysis buffer (Millipore) containing protease inhibitor cocktails (Roche Diagnostics, Mannheim, Germany). Equal amounts of protein were loaded on SDS-polyacrylamide electrophoresis gels and transferred onto polyvinylidene difluoride membranes after electrophoresis. The primary antibodies used were as follows: mouse anti-Synaptophysin (1:1000, MAB5258A4, Merck), mouse anti-PSD 95 (1:1000, MA1-046, Invitrogen), rabbit anti-C3aR (1:1000, sc-133172, Santa Cruz), mouse anti-Apolipoprotein E4 (1:1000, M067-3, MBL Life Science), rabbit anti-TPK1 (1:1000, ab230263, Abcam), rabbit anti-APP (1:1000, ab32136, Abcam), rabbit anti-BACE1 (1:1000, 5606, CST), mouse anti-Tau5 (1:1000, AHB0042, Invitrogen), rabbit anti-pTau (Thr181) (1:1000, 12885, CST), rabbit anti-pTau (Ser396) (1:1000, 9632, CST), mouse anti-pTau (Ser202, Thr205) (1:1000, MN1020, Invitrogen), rabbit anti-β-actin (1:2000, 4970, CST), rabbit anti-tubulin (1:2000, 2146, CST). The membranes were incubated overnight with primary antibodies, followed by incubation with horseradish peroxidase-conjugated anti-rabbit, anti-mouse, or anti-rat secondary antibodies. Blots were processed using Image Lab-6.0 software and Image J.

### Immunohistochemistry

After being anesthetized, mice were transcardially perfused with ice-cold PBS and fixed for 24 h in 4% PFA/PBS at 4 °C. The brains were taken out and then transferred to 30% sucrose for 2 days. Immunohistochemical staining was conducted on 20 μm frozen coronal sections. The following primary antibodies were used: rat anti-Iba1 (1:500, ab283346, Abcam), goat anti-Iba1 (1:200, ab5076, Abcam), goat anti-GFAP (1:200, ab53554, Abcam), mouse anti-NeuN (1:200, MAB377, Millipore), rabbit anti-CD68 (1:200, ab283654, Abcam), sheep anti-trem2 (1:200, AF1729, R&D), rabbit anti-4G8 (1:200,800702, Bio Legend), rabbit anti-TSPO (1:2000, ab109497, Abcam), rat anti-C3 (1:200, ab11862, Abcam), and rabbit anti-C1q (1:200, ab182451,abcam).

The secondary antibodies used were as follows: Alexa Fluor 594-conjugated donkey anti-rabbit IgG (1:1000, A-21207), Alexa Fluor 488-conjugated goat anti-rabbit IgG (1:1000, A-11008), Alexa Fluor 594-conjugated donkey anti-goat IgG(1:1000, A-11058), Alexa Fluor 594-conjugated donkey anti-rat IgG (1:1000, A-21209), Alexa Fluor 594-conjugated goat anti-mouse IgG (1:1000, A-11005), Alexa Fluor 488-conjugated donkey anti-goat IgG (1:1000, A-21447), and Alexa Fluor 594-conjugated anti-sheep IgG (1:1000, A11016, all from Invitrogen, Waltham, MA). DNA was stained with Hoechst 33342 (1:10,000, H3570, Invitrogen). Staining was visualized by an Olympus FV 3000 laser-scanning confocal microscope and a Nikon A1R confocal microscope.

### Bulk RNA-seq

Bulk RNA-seq was performed by Shanghai Majorbio Bio-Pharm Technology. mRNAs were enriched from total RNA samples using poly-A pulldown, and cDNA libraries were prepared using Illumina Truseq chemistry, followed by sequencing with Illumina NovaSeq 6000. The sequencing data of transcriptome based on the Illumina sequencing platform were scientifically counted and strictly controlled to reflect the library quality and sequencing quality of this experiment. The QC data was compared with the reference genome constructed by HISAT to obtain mapped data. After the transcript was quantitatively analyzed and its Read Counts were obtained, the differential expression analysis between groups was carried out to identify the differentially expressed genes between different groups, and then the function of the differentially expressed genes was studied.

### SB290157 preparation and treatment

SB290157 (MedChemexpress) solution was prepared and treated as in previous studies [19, 20]. Briefly, SB290157 was dissolved in DMSO and then diluted by PBS up to 100 μg/ml before usage. The *Tpk*-cKO mice with humanized APOE4 knock-in (AE-cKO mice) were randomly assigned to the drug treatment group or the PBS control group. After 4 weeks after tamoxifen induction, the AE-cKO mice were intraperitoneally injected with SB290157 or PBS containing equal DMSO three times a week at a dose of 1 mg/kg according to the mouse body weight for six weeks.

Behavioral tests were performed in the 8^th^ week after tamoxifen induction and all of the mice were anesthetized and dissected in the 10^th^ week after tamoxifen induction (Fig. 7a).

### Statistical analysis

GraphPad Prism 9.5 (GraphPad software, GraphPad Software, Inc., San Diego, California, USA) was used for statistical analyses. One-way ANOVA for multiple comparisons with appropriate Tukey’s or Dunnett’s multiple comparison tests was used to identify statistical differences. The results were represented as the mean ± SEM. ‘N’ refers to the number of animals, unless otherwise indicated. All conditions that were statistically different from its control are indicated, where * represents P < 0.05.

## Results

### *APOE4* aggravates behavior dysfunction of TPK cKO mice

To unravel the roles of *APOE4*, we crossbreed the *Tpk-*cKO mice with mice that were knocked in human *APOE4* (AE mice) and generated the AE-cKO mice (Fig. 1a). All kinds of genotype mice were intraperitoneally injected with tamoxifen to induce Cre expression or as control at 10 weeks-old adult mice, then the behavior and pathologies were examined at 18 and 20 weeks old respectively (Fig. 1b). Western blot studies showed that human APOE4 protein was well expressed in brains of the AE and AE-cKO mice, and TPK protein was significantly reduced in the *tpk-*cKO and AE-cKO mice, indicating that the mouse models were successfully established (Fig1. c, d).

**Fig. 1:**
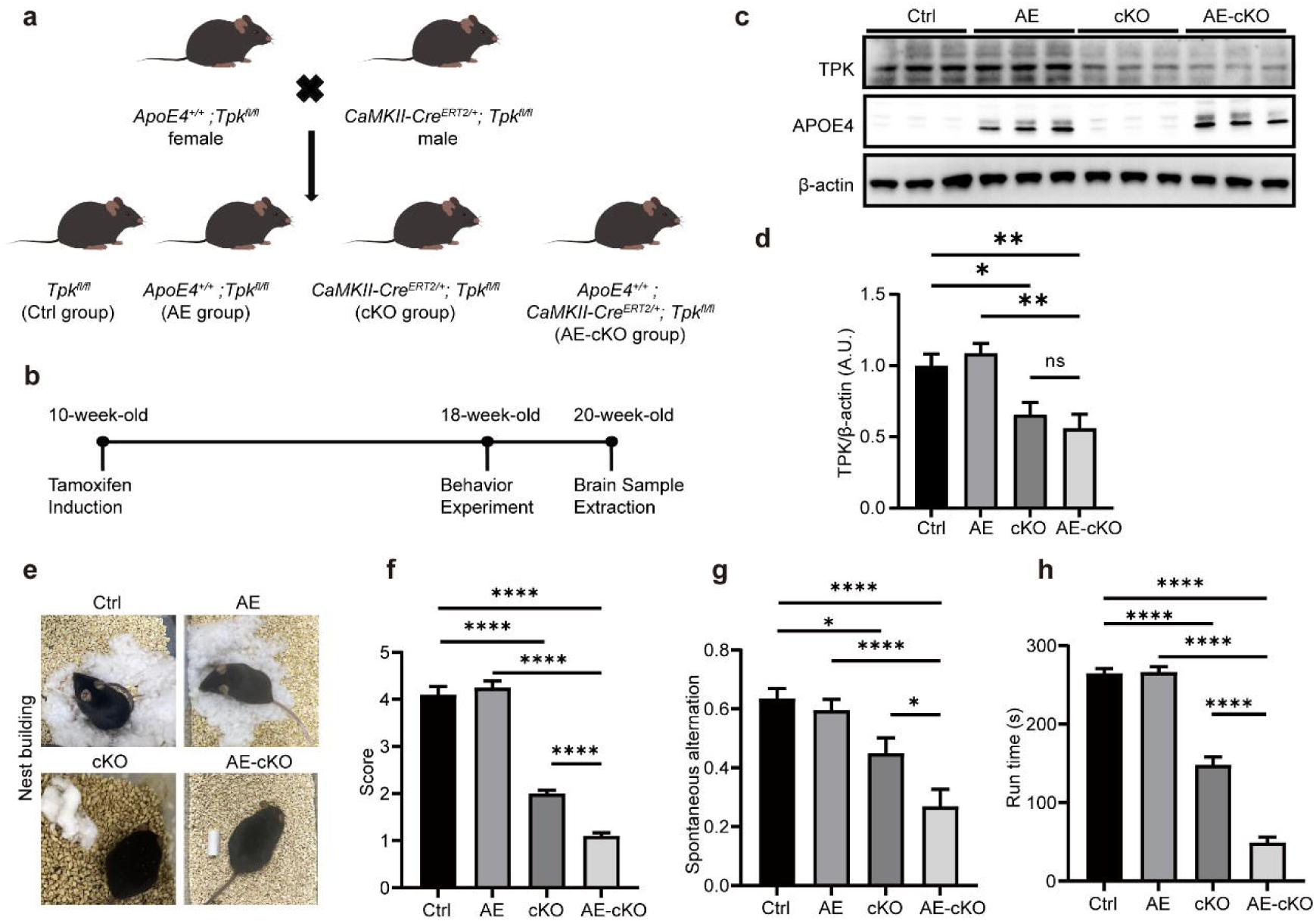
APOE4 exacerbates behavior abnormalities of mice induced by neuronal *Tpk* deletion. **a.** Establishment and grouping of mouse models. **b.** The diagram of experimental workflow. **c, d.** Representative images of western blotting (**c**) and quantification (**d**) of TPK protein level in the cortices of mice. The results showed that the humanized *APOE4* knock-in did not significantly affect the level of brain TPK protein of mice with or without *Tpk* deletion in neurons. N = 5 mice per group. **e, f.** Representative images and scores of nest building. The results showed that the humanized *APOE4* knock-in significantly exacerbated the abilities of nest building of the cKO mice, but did not change those of the mice without *Tpk* deletion. N = 20 per group. **g.** Y-maze test. The results showed that the humanized *APOE4* knock-in significantly aggravated the behavior abnormality of the cKO mice but did not affect the behavior of the mice without *Tpk* deletion. N = 20 in Ctrl and AE groups. N = 18 in cKO and AE-cKO groups. **h.** Rotarod test. The results showed that APOE4 significantly deteriorated the reduction of run time on the rotarod of the cKO mice, but did not alter the run time on the rotarod of the mice without *Tpk* deletion. N = 20 per group. The data represent mean ± SEM. ns: not significant, \**p*<0.05, \*\**p*<0.01, \*\*\*\**p*<0.0001, one-way ANOVA followed by Tukey’s *post hoc* test.

To determine whether the presence of human *APOE4* isoforms affects TPK expression, we compared the levels of TPK protein in all of the four groups of mice. There were no significant changes in the levels of brain TPK protein between the AE and control groups. Similarly, *APOE4* doesn’t affect the levels of brain TPK in the AE-cKO mice as compared with that in the *Tpk-*cKO mice (Fig. 1c, d). Those results indicate that APOE4 not only doesn’t directly affect brain TPK expression.

Consistent with our previous study ^15^, the *Tpk-*cKO mice showed significant impairment of nest building, spatial memory, and motor ability assessed by nest building, Y-maze, and rotarod behavior assays. Here, we further found that the AE-cKO mice exhibited more severe abnormalities in nest building, Y-maze, and rotarod tests (Fig. 1e-h). However, there were not significant changes between Ctrl and AE groups. The results indicate that *APOE4* significantly accelerated the progression of behavior dysfunction in the *Tpk-*cKO mice, but did not affect the behavioral results of control mice.

### Aggravated synapse loss in the AE-cKO mice

The pathological basis of the exacerbated behavior dysfunction in the AE-cKO mice was further investigated. We found that *APOE4* worsened the loss of brain synapses induced by neuronal *Tpk* deletion, presented by further reduction of synaptic proteins PSD95 and Synaptophysin in brains of the AE-cKO mice as compared with those in the *Tpk-*cKO mice. However, *APOE4* did not affect the loss of brain neurons (Fig. 2a-d). Also, there were no significant changes in the levels of synaptic proteins and the number of NeuN-positive neurons in the AE mice compared to the control mice (Fig. 2e, f). Those results suggest that *APOE4* specifically aggravates synapse loss, but does not affect neuron loss, in the brains of the *Tpk*-cKO mice.

**Fig. 2:**
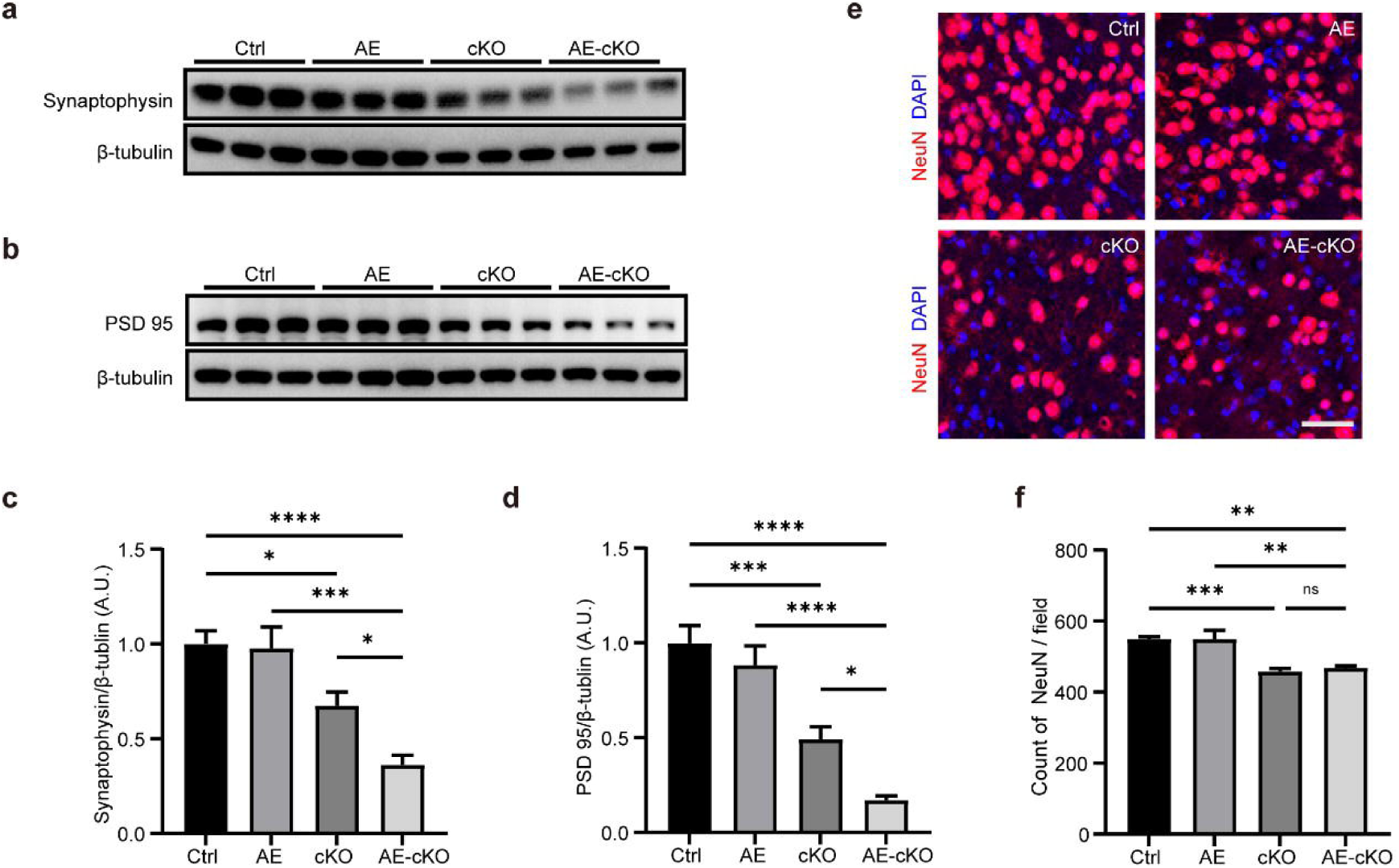
APOE4 deteriorates loss of brain synapses, but does not affect loss of brain neurons of mice induced by neuronal *Tpk* deletion. **a-d.** Representative images of western blotting (**a, b**) and quantification (**c, d**) of synaptophysin and PSD95 in the cortices of mice. The results showed that the humanized *APOE4* knock-in significantly exacerbated the reduction of synaptophysin (**a, c**) and PSD95 (**b, d**) in cortices of the cKO mice, but did not alter those in the mice without *Tpk* ablation. N = 6 mice per group. **e, f.** Representative immunostaining images of NeuN (red, **e**) and quantifications (**f**) in the cortices of mice. The results showed that the humanized *APOE4* knock-in did not significantly change the number of cortical neurons in the mice with or without *Tpk* depletion in neurons. N = 5 mice per group. The data is represented by mean ± SEM. ns: not significant, \**p*<0.05, \*\**p*<0.01, \*\*\**p*<0.001, \*\*\*\**p*<0.0001, one-way ANOVA followed by Tukey’s *post hoc* test. Scale bars, 25 μm.

### Altered transcriptional signatures in AE-cKO mice

To further investigate the mechanism of synapse loss observed in the AE-cKO mice, we performed RNA sequencing analysis. RNA sequencing of cortical samples in the 10^th^ week after tamoxifen treatment showed that 339 genes were significantly upregulated and 287 genes were downregulated in the AE-cKO mice compared to the *Tpk-*cKO mice (Fig. 3a, b). Based on analyses of gene ontology (GO), gene set enrichment analysis (GSEA), and the Kyoto Encyclopedia of Genes and Genomes (KEGG) pathway, significant enrichments were found in genes associated with the signal pathways that regulate phagocytosis, glutamatergic synapse, complement-dependent synaptic pruning, complement complex (such as C1, C3, and C5), etc. (Fig. 3c-e).

**Fig. 3:**
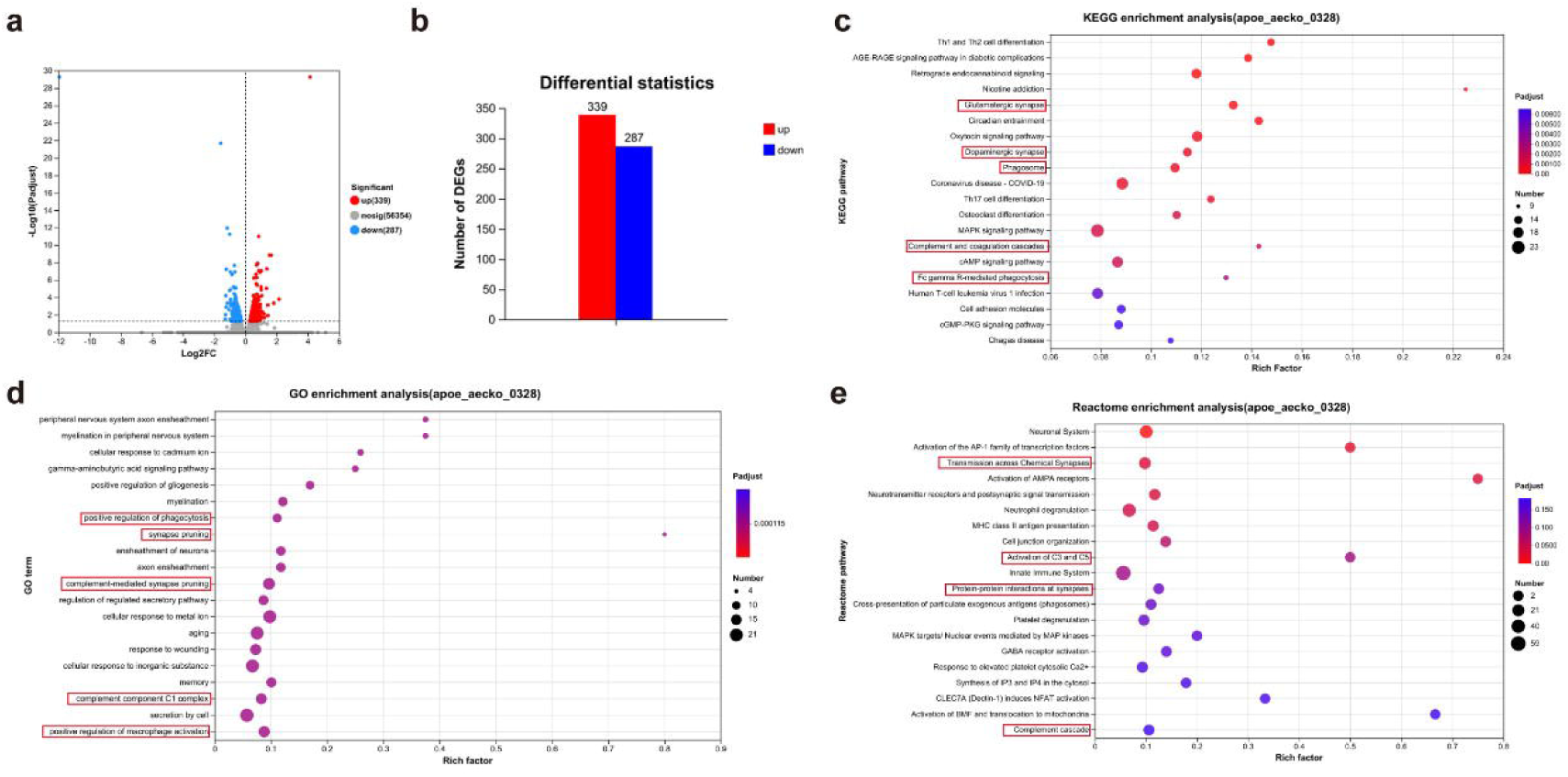
APOE4 regulates the expression of genes associated with microglia and the complement-related pathway in mice induced by neuronal *Tpk* deletion. Identification of differentially expressed genes and functional gene enrichment analyses. **a.** Volcano plot. **b.** The number of DEGs. **c.** GO analyses of DEGs. **d.** KEGG pathway analysis of DEGs. **e.** Reactome pathway analysis of DEGs.

### The phagocytosis of synapses by microglia was significantly intensified in the AE-cKO mice

Numerous studies have highlighted the important role of microglia in the phagocytosis of synapses [22]. Therefore, combined with the results of our transcriptomic analysis, we further examined the effect of *APOE4* on synapse phagocytosis of microglia in the *Tpk*-cKO mice. Immunohistochemistry staining of Iba1 showed that microglia activation was more significant in the AE-cKO mice compared to the *Tpk-*cKO mice (Fig. 4a, b). The results were further validated by staining with TSPO antibody, an activated microglia marker, which TSPO-positive microglia were significantly enhanced in the AE-cKO mice compared to the *Tpk-*cKO mice (Fig. 4c, d). Functionally, the activated microglia were accelerated to translate to the status of disease-associated and phagocytosis, represented by significantly enhanced expression in CD68 and TREM2 of microglia in the AE-cKO mice as compared with those in the *Tpk-*cKO mice (Fig. 4e-h). Furthermore, co-staining of Iba1 and PSD95 antibodies showed that the reactive microglia engulfed post-synapse protein PSD95 (Fig. 4i). Taken together, those results demonstrate that *APOE4* promotes the activation and transition of microglia to the status of disease-associated and synaptic phagocytosis that mediate synapse elimination.

**Fig. 4:**
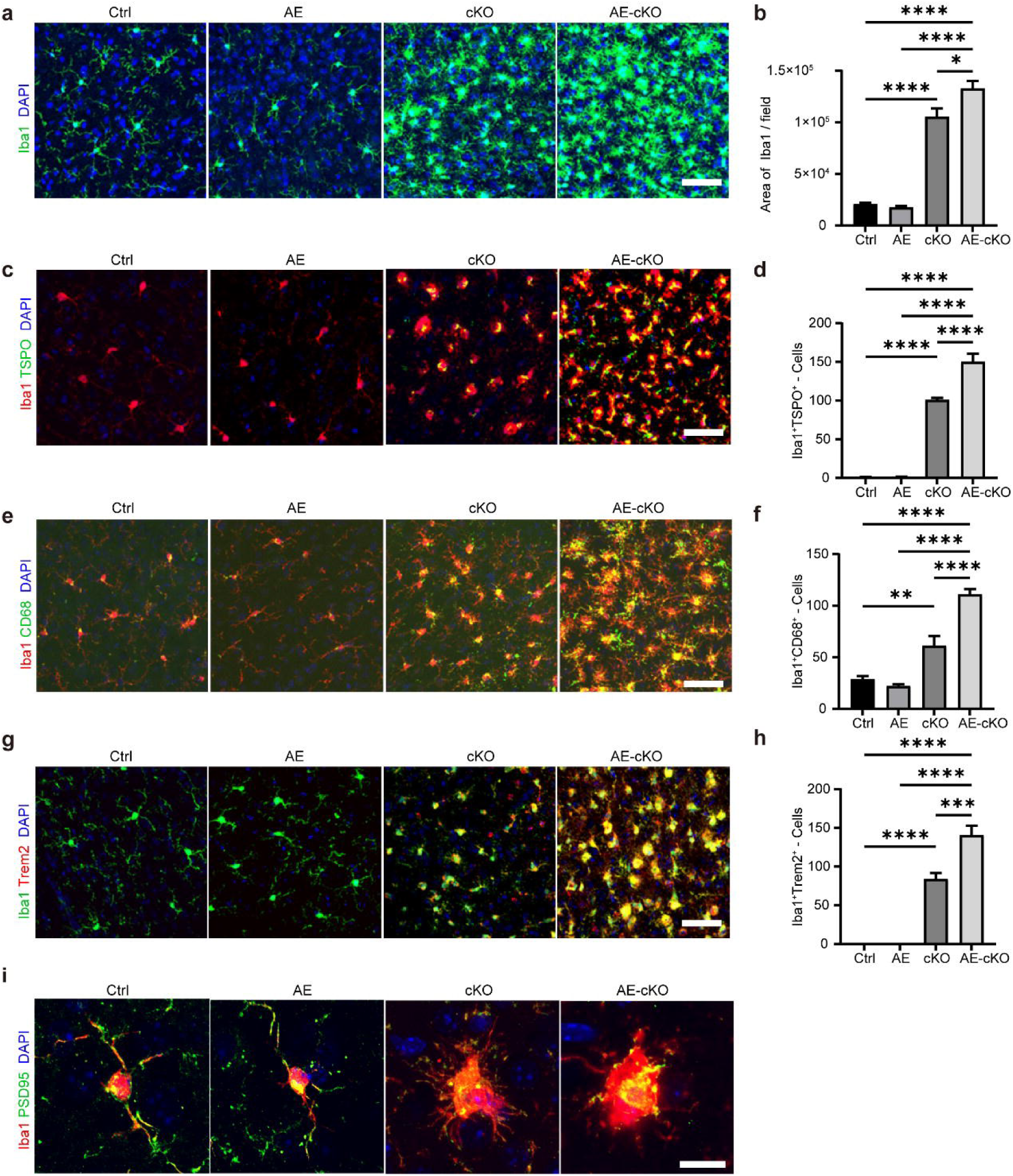
APOE4 aggravates microglial activation in mice induced by neuronal *Tpk* deletion. **a, b.** Representative images of Iba1 (green) immunostaining (**a**) and quantification (**b**) in the cortices of mice. The results showed that the humanized *APOE4* knock-in significantly worsened the activation of microglia in the cortices induced by neuronal *Tpk* ablation, but did not affect those in the mice without *Tpk* knockout. **c, d.** Representative images of Iba1 (red) and TSPO (green) immunostaining (**c**) and quantification (**d**) in the cortices of mice. The results showed that the humanized *APOE4* knock-in significantly upregulated the enhancement of co-labeled cells by Iba1 and TSPO antibodies in the cKO mice, but did not affect those in mice without *Tpk* knockout. **e, f.** Representative images of Iba1 (red) and CD68 (green) immunostaining (**e**) and quantification (**f**) in the cortices of mice. The results showed that the humanized *APOE4* knock-in significantly exacerbated the elevation of co-labeled cells by Iba1 and CD68 antibodies in the cKO mice, but did affect those in mice without *Tpk* knockout. **g, h.** Representative images of Iba1 (green) and Trem2 (red) immunostaining (**g**) and quantification (**h**) in the cortices of mice. The results showed that the humanized *APOE4* knock-in significantly aggravated the enhancement of co-labeled cells by Iba1 and Trem2 antibodies in the cKO mice but did not affect those in mice without *Tpk* knockout. N = 5 mice per group. The data represent mean ± SEM. \**p*<0.05, \*\**p*<0.01, \*\*\**p*<0.001, \*\*\*\**p*<0.0001, one-way ANOVA followed by Tukey’s *post hoc* test. Scale bars, 25 μm. **i** Representative images of immunostaining of Iba1 (red) and PSD95 (green) i n the cortices of mice. The results showed that microglia-mediated synaptic phagocytosis presented by PSD95 in the microglia. Scale bars, 10 μ

### *APOE4* enhanced synaptic pruning *via* microglia-mediated complement-dependent pathway in the AE-cKO mice

Previous studies have shown that microglial engulfment and elimination of synapses are mediated by the classical complement cascade [22, 23]. In this cascade, the initiating molecule C1q and its downstream C3 localize to synapses and provoke the engulfment and elimination of synapses of microglia with that express C3 receptor (C3R) [22]. Here we found that C1q and co-localization of C1q with Iba1 was significantly enhanced in the AE-cKO mice compared to the *Tpk*-cKO mice (Fig. 5a, b). C3 was also significantly increased in the AE-cKO mice as compared with that in the *Tpk*-cKO mice (Fig. 5c, d). Immunoblot analysis of C3R also showed that *APOE4* significantly enhanced the level of C3R protein in the AE-cKO mice as compared with that in the *Tpk*-cKO mice (Fig. 5e, f). Those results suggest that APOE4 contributes to synapse loss by increasing microglia-mediated complement-dependent synaptic pruning in the AE-cKO mice.

**Fig. 5:**
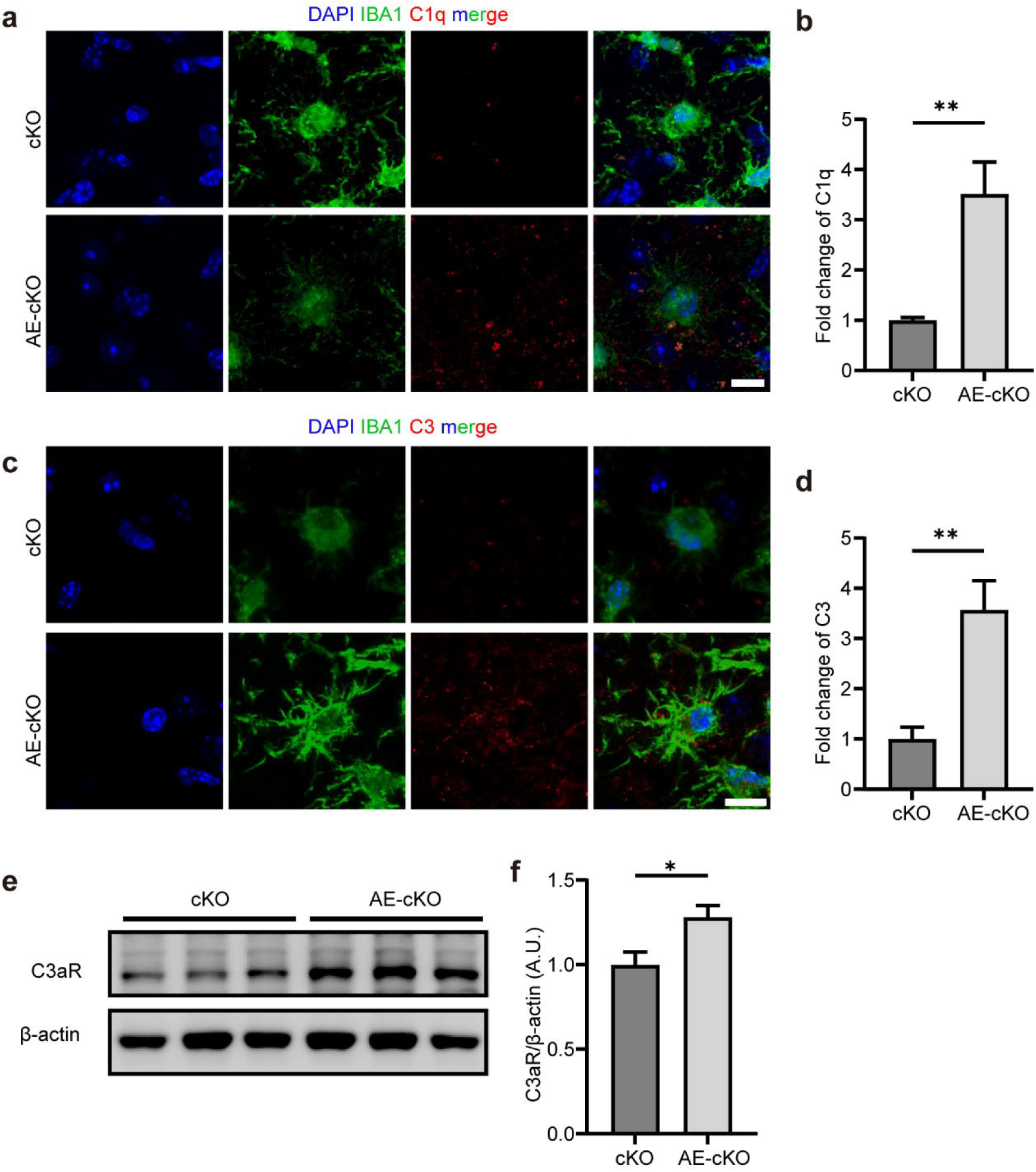
APOE4 enhances complement-mediated synaptic pruning in mice induced by neuronal *Tpk* deletion. **a, b.** Representative images of Iba1 (green) and C1q (red) immunostaining (**a**) and quantification (**b**) in the cortices of mice. The results showed that the humanized *APOE4* knock-in significantly aggravated the increase of c1q expression induced by neuronal *Tpk* deletion N = 5 mice per group. **c, d.** Representative images of Iba1 (green) and C3 (red) immunostaining (**c**) and quantification (**d**) in the cortices of mice. The results showed that the humanized *APOE4* knock-in significantly worsened the increase of C3 expression induced by neuronal *Tpk* deletion. N = 5 mice per group. **e, f.** Representative images of Western blotting (**e**) and quantification (**f**) of C3aR protein in the cortices of mice. The results showed that the humanized ApoE4 knock-in significantly accelerated the increase of C3aR protein induced by neuronal *Tpk* deletion. N = 6 mice per group. The data represent mean ± SEM. \**p*<0.05, \*\**p*<0.01, unpaired *t*-test. Scale bars, 10 μ

### *APOE4* does not affect Aβ pathologies, Tau hypophosphorylation, and astrocyte activation induced by neuronal *Tpk* deletion

The effects of *APOE4* on Aβ pathologies, Tau hypophosphorylation, and astrocyte activation of mice induced by neuronal *Tpk* deletion were also investigated. The previous studies have shown that APOE4 contributes to Aβ deposition by direct interaction with Aβ [7, 9]. However, here we didn’t find that *APOE4* further promoted the enhancement in the levels of APP, BACE1, CTFα, and CTFβ proteins in the AE-cKO mice as compared with those in the *Tpk*-cKO mice (Fig. 6a-f). Consistently, the levels of soluble Aβ40 and Aβ42 and the ratio of Aβ42/40 were not significantly changed in the AE-cKO mice as compared with those in the *Tpk*-cKO mice (Fig. 6g-i). Staining of Aβ plaques using 4G8 antibody also showed that the Aβ plaques in the AE-cKO mice was not significantly affected as compared with that in the *Tpk*-cKO mice (Fig. 6j, k). The levels of p-Tau and total Tau were detected by Western blot. The results showed that the levels of Tau phosphorylation and the ratios of p-Tau to total Tau at S202, T205, S396, and T181 sites was not significantly changed in the AE-cKO mice as compared with those in the *Tpk*-cKO mice (Fig. 6l-o). GFAP staining showed that *APOE4* did not affect the activation of astrocytes caused by neuronal *Tpk* deletion(Fig. 6p, q).

**Fig. 6:**
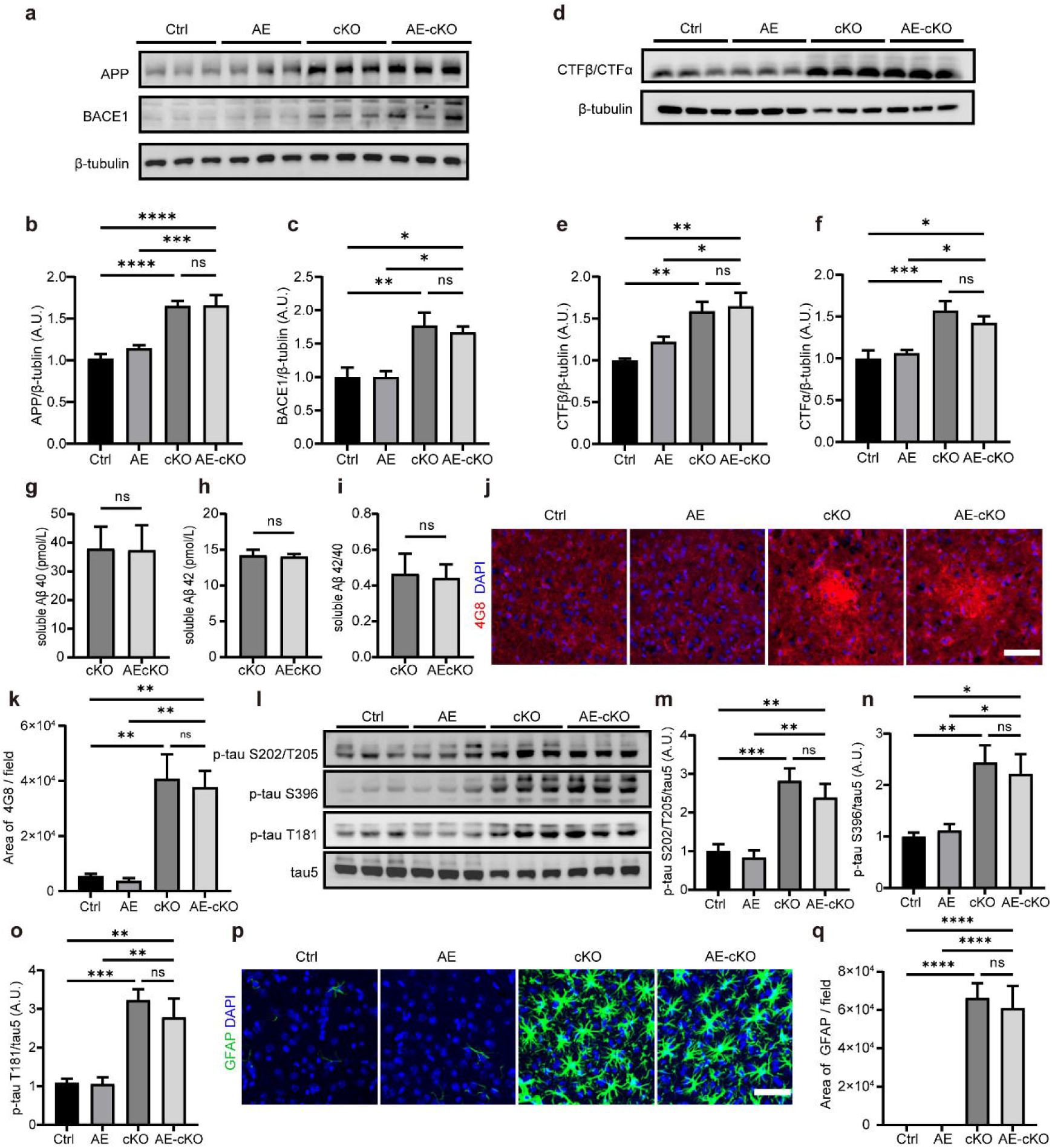
APOE4 does not affect astrocyte activation and Aβ plaques of mice induced by neuronal *Tpk* deletion. **a-f.** Representative images of western blotting (**a, d**) and quantification (**b, c, e, f**) of APP, BACE1 C99, and C83 in the cortices of mice. The results showed that the humanized *APOE4* knock-in did not exacerbate the increase of APP (**a, b**), BACE1 (**a, c**), and CTFβ/CTFα (**d-f**) in the cortices of the *Tpk*-cKO mice, and did not alter the increase of APP (**a, b**), BACE1 (**a, c**) and CTFβ/CTFα (**d-f**) in mice without *Tpk* ablation. N = 6 mice per group. **g-i.** Representative images of ELISA quantification (**g-i**) of soluble Aβ40, Aβ42, and 40/42 in the cortices of mice. The results showed that *APOE4* overexpression did not affect the expression of soluble Aβ40 (**g**), Aβ42(**h**), and 40/42 (**i**) in the cortex of the *Tpk*-cKO mice. N=5 **j, k.** Representative images of 4G8 (red) immunostaining (**j**) and quantification (**k**) in the cortices of mice. The results showed that the humanized *APOE4* knock-in did not affect the number of Aβ plaques in the cortices of mice with or without *Tpk* deletion. **l-o.** Representative images of western blotting (**l**) and quantification (**m-o**) of p-tau S202/T205, p-tau S396, p-tau T181 and tau 5 in the cortices of mice. The results showed that the humanized *APOE4* knock-in did not exacerbate the increase of p-tau/tau at S202/T205, S396, and T181 in the corticies of the *Tpk*-cKO mice, and did not alter the increase of ratios of p-tau/tau at S202/T205, S396, and T181 sites in mice without *Tpk* ablation. N = 6 mice per group. **p, q.** Representative images of GFAP (green) immunostaining (**p**) and quantification (**q**) in the cortices of mice. The results showed that the humanized *APOE4* knock-in did not exacerbate the activation of cortical astrocytes in mice with or without *Tpk* deletion in neurons. N = 5 mice per group. The data represent mean ± SEM. ns: not significant, \**p*<0.05, \*\**p*<0.01, \*\*\**p*<0.001, \*\*\*\**p*<0.0001, one-way ANOVA followed by Tukey’s *post hoc* test. Scale bars, 25 μ

### C3aR antagonist significantly repressed microglia phagocytosis and ameliorated synapse loss in the AE-cKO mice

SB290157, an antagonist of C3aR [19, 20], was used to further evaluate the role of *APOE4* in enhancing microglia-mediated complement-dependent synaptic pruning in the AE-cKO mice. After 4 weeks of tamoxifen induction, the AE-cKO mice were intraperitoneally injected with SB290157 (1mg/kg) or control vehicle, and then the behavior and pathologies were examined at 18 and 20 weeks old respectively (Fig. 7a). No significant differences in behavior scores were found in nest building, Y-maze, and rotarod assays between the AE-cKO mice with and without SB290157 treatment (Fig. 7b-d). However, Iba-1 immunostaining showed that the area of microglia was significantly reduced by SB290157 treatment, but the microglia number was not obviously changed (Fig. 7e-g). Those results show that SB290157 inhibits the amoeboid transformation of microglia induced by neuronal *Tpk* deficiency, suggesting that microglia engulfment ability is repressed. Further, the levels of CD86, TSPO, and Trem2 proteins in the AE-cKO mice with SB290157 treatment were significantly decreased as compared with those in the AE-cKO mice without SB290157 treatment (Fig. 7h-m). Furthermore, the levels of PSD95 and Synaptophysin proteins were significantly ameliorated in the AE-cKO mice with SB290157 treatment as compared with those with the vehicle (Fig. 7n-p). Taken together, these results suggest that *APOE4* promotes the synapse lost in the AE-cKO mice through microglia-mediated complement-dependent synaptic pruning.

**Fig. 7:**
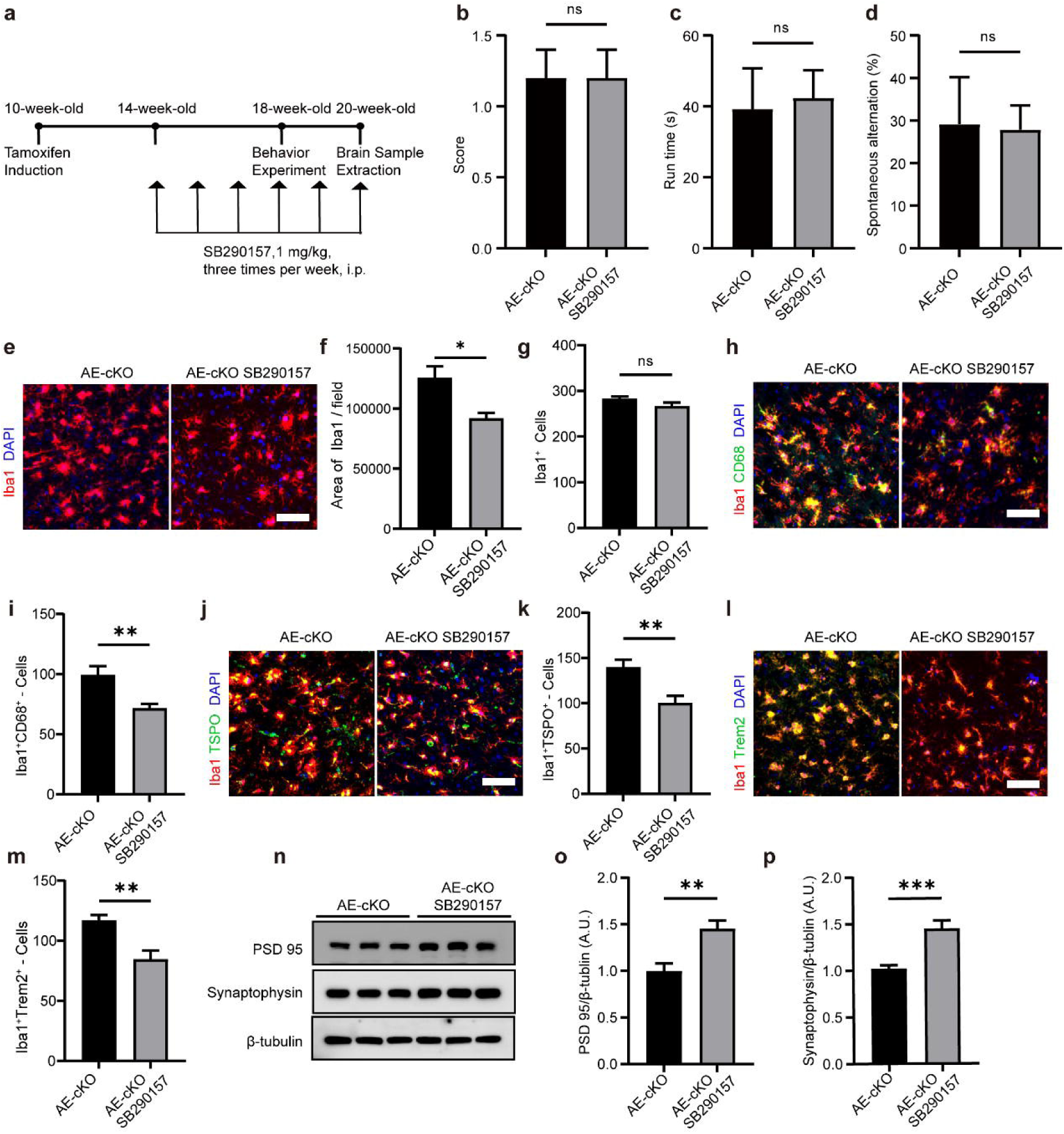
C3aR antagonist SB290157 significantly represses the exacerbation of microglia activation and synapse loss induced by APOE4, but not improve behavior abnormalites due to neuronal *Tpk* deletion. **a.** Flow chart of injection of SB290157. The injection of SB290157 started from the 4 weeks after tamoxifen induction, the frequency of injections is 3 times per week. **b.** Scores of nest building. The results showed that the injection of SB290157 did not alleviate the abilities of nest building of the AE-cKO mice. N = 5 per group. **c.** Rotarod test. The results showed that the injection of SB290157 did not alleviate the run time on the rotarod of the AE-cKO mice. N = 5 per group. **d.**Y-maze test. The results showed that the injection of SB290157 did not alleviate the behavior abnormality of the AE-cKO mice. N = 5 per group. **e-g.** Representative images of Iba1 (red) immunostaining (**e**) and quantification (**f,g**) in the cortices of mice. The results showed that the injection of SB290157 significantly decreased the activation of microglia in the cortices of the AE-cKO mice, but did not affect the count of Iba. **h, i.** Representative images of Iba1 (red) and CD68 (green) immunostaining (**h**) and quantification (**i**) in the cortices of mice. The results showed that the injection of SB290157 significantly decreased the elevation of co-labeled cells stained by Iba1 and CD68 antibodies in the AE-cKO mice. **j, k.** Representative images of Iba1 (red) and TSPO (green) immunostaining (**j**) and quantification (**k**) in the cortices of mice. The results showed that the injection of SB290157 significantly downregulated the enhancement of co-labeled cells stained by Iba1 and TSPO antibodies in the AE-cKO mice. **l, m.** Representative images of Iba1 (red) and Trem2 (green) immunostaining (**l**) and quantification (**m**) in the cortices of mice. The results showed that the humanized *APOE4* knock-in significantly decreased the elevation of co-labeled cells stained by Iba1 and Trem2 antibodies in the AE-cKO mice. N = 5 mice per group. **n-p.** Representative images of western blotting (**n**) and quantification (**o, p**) of synaptophysin and PSD95 in the cortices of mice. The results showed that the injection of SB290157 significantly alleviated the reduction of synaptophysin (**n, p**) and PSD95 (**n, o**) in the cortices of the AE-cKO mice. N = 6 mice per group. The data represent mean ± SEM. \**p*<0.05, \*\**p*<0.01, \*\*\**p*<0.001, \*\*\*\**p*<0.0001, one-way ANOVA followed by Tukey’s *post hoc* test. Scale bars, 25 μm.

## Discussion

In the current studies, we found that APOE4 significantly exacerbates the loss of brain synapses and hence behavioral abnormalities induced by *Tpk* deletion in neurons, but doesn’t directly affect brain TPK expression (Fig. 1). The results are similar to those in the previous studies, which APOE4 leads to significant deteriorations of Aβ and Tau pathologies in Aβ and/or tau transgenic mouse models, but does not triggers Aβ and Tau pathological alterations by itself in the wild-type and/or control mice [6–11]. Because APOE4 does not simultaneously exacerbate the loss of neurons in the *Tpk*-cKO mice (Fig. 2), the results suggest that the deterioration of synaptic loss occurs at the synapse itself. The unconventional pruning of synapses mediated by the overactivated complement pathway of anomaly microglia is considered to be critical for the synaptic loss of neurodegenerative diseases [24, 25].

Therefore, the role of microglial transition and the associated complement pathway was further investigated. The results showed that APOE4 significantly promotes the transition of activated microglia into phagocytosis status and leads to the overactivation of corresponding complement pathways (Fig. 3 & 4), which results in the pronounced pruning of synapses involved in microglia in the AE-cKO mice (Fig. 5). SB290157, a specific C3aR antagonist, significantly ameliorated synapse loss in the AE-cKO mice further supported this notion (Fig. 7). The phenomenon is independent of the alterations of Aβ deposition, Tau pathologies, and astrocyte activation (Fig. 6).

To date, the molecular pathobiology underlying the effect of APOE4 on the onset and progression of AD remains puzzling. Neuroinflammation is suggested to be a major factor in the mechanism by which APOE4 influences the disease [12]. Microgliosis promoted Tau-mediated neurodegeneration in the presence of APOE4 [11]. On the other side, the ablation of microglia prevents APOE4-mediated neurodegeneration and also reduces Tau phosphorylation in mice with Tau pathological alteration [26]. In addition, perturbed lipid and glucose metabolism may contribute to APOE4-associated dysfunction of microglia [12, 13]. Pharmacological and genetic methods promoting lipid metabolism in microglia strongly attenuated APOE4-mediated neurodegeneration in the P301S/APOE4 mice [27]. These results are consistent with our current studies, indicating that the pathological transition of microglia is involved in the mechanism of worsening pathophysiological alterations of AD caused by APOE4.

Microglia play an important role not only in maintaining the homeostasis of normal synapse plasticity [28] but also in leading to synaptic dysfunction of neurological diseases [29, 30]. Further, APOE4 significantly enhances C1q accumulation in the brain as compared with APOE3 allele, while APOE2 decreases C1q level [31]. Therefore, C1q accumulation may be involved in the mechanism of APOE4 eliminating synapses mediated by activated microglia. In this study, we for the first time found that APOE4 promoted C1q, C3, and C3aR expression and activated complement-dependent synaptic pruning in the mice. Further, we found that C3aR antagonist significantly repressed microglia phagocytosis and ameliorated synapse loss in the AE-cKO mice. These results show that the APOE4-mediated worsening of synapse loss caused by neuronal *Tpk* deficiency is involved in microglia-associated complement pathway, promoting the understanding on the mechanism by which APOE4 exacerbates AD progression.

However, our studies also found that C3aR antagonist SB290157 did not significantly improve the abnormalities in behavior tests of the AE-cKO mice, suggesting that only SB290157 treatment is insufficient. It may be attributed to the following two reasons: First, SB290157 can not completely inhibit the activation of the microglial complement pathway. As in Fig 2, the protein levels of PSD95 and Synaptophysin in the *Tpk*-cKO mice were 2 and 1.8 times as compared with those in the AE-cKO mice, while SB290157 treatment only enhanced the levels of PSD95 and Synaptophysin proteins to 1.5 times in the AE-cKO (Fig 7). Second, other mechanism(s) may contribute to the APOE-mediated aggravation of synapse loss induced by neuronal *Tpk* deletion, leading to insufficiently rescue abnormalities of behavior tests in the mice. The status and underlying mechanism of microglia overactivation caused by APOE should continue to be explored in future studies.

In summary, APOE4 aggravates synaptic impairment and hence behavior abnormalities induced by *Tpk* deletion in neurons. It may be involved in the disease-associated transition of microglia via the complement pathway due to C1q accumulation in the brain. Combined with the results from our current study and the previous studies [15, 16, 32–34], a preliminary conclusion can be drawn, which APOE4 exacerbates multi-pathophysiological cascades of AD that have already occurred, just like adding fuel to the fire. Other risk factors of AD, such as diabetes, may directly induce multiple pathophysiological impairments of AD [16], which is like a spark that directly ignites a fire and is different from the role of APOE4 in the pathogenesis of AD. Elucidating the specific link(s) and mechanism(s) by which risk factors act on the pathogenesis of AD is an essential foundation for a comprehensive understanding of the occurrence and development of the disease. Our current studies promote the understanding of AD pathogenesis.

## Acknowledgements

This study was supported by grants from the Shanghai Municipal Science and Technology Major Project, and the National Natural Science Foundation of China (81870822, 91332201, 81901081, 81600930,

## Author contributions

Pengyue Du, Boru Jin, and Zijie Wang were responsible for pathological and behavior experiments. Boru Jin was responsible for transcriptional RNA sequencing. Shaoming Sang and Binqiao Zhao were responsible for the verification of the results. Shaoming Sang, Binqiao Zhao, and Chunjiu Zhong designed the study and wrote the manuscript.

## Competing interests

Chunjiu Zhong, one of the corresponding authors, holds shares of Shanghai Raising Pharmaceutical Co., Ltd., which is dedicated to developing drugs for the prevention and treatment of AD. The other authors declare that they have no competing interests.

